# Bio-electricity production from Fibroblasts and their cultivation media

**DOI:** 10.1101/2024.01.04.574251

**Authors:** Yaniv Shlosberg, Ayelet Lesman

## Abstract

The recent understanding of the critical future damage that might happen on earth by climate change has urged scientists to initiate new creative ideas for clean energy technologies that will reduce carbon emissions. A promising approach is the utilization of living cells as electron donors in bio-electrochemical cells (BECs). This concept has been intensively studied for micro-organisms such as non-photosynthetic bacteria, cyanobacteria, and microalgae, but not for mammalian cells. In this work, we report for the first-time integrating live fibroblast cells in a BEC to produce electrical current that is about 3 times higher than intact micro-organisms. Furthermore, we apply 2D-fluorescence and electrochemical measurements to show that like in micro-organisms based BECs, NADH and flavins play a role in the electron mediation between the cells and the anode. Finally, we show that the cultivation medium of fibroblasts also consists of redox species that may produce dark and photocurrent.

## Introduction

In recent years, the climate crisis has driven scientist around the world to develop clean energy approaches that can supply the increasing global energy demand and reduce carbon emissions replacing the traditional fossil fuel-based technologies. A unique method for electricity generation is the use of living cells as electron donors or acceptors in bio-electrochemical cells (BECs). This concept was first reported more than 100 years ago by Potter et al.^1^ which integrated bacteria with an electrochemical cell inventing the microbial fuel cells (MFCs). Over the years, the power output of MFCs was significantly improved by using bacterial species with a high exo-electrogentic activity such as *shewanella oneidensis*^2–5^ and *Geobacter* sp.^6–8^ that can conduct a direct electron transfer to the anode of the BEC by conductive protein complexes such as *pili* and metal respiratory complexes^9–11^. The electron transfer may also be conducted by diffusive redox species that can be released from the cells such as NADH, flavins, vitamins and phenazines^2,12–14^. Rather than endogenous bio-electron mediators, the electron transfer can be enhanced by adding exogenous electron mediators such as potassium ferricyanide^15^. To maintain MFCs, there is a need to supply carbon sources for the bacteria to survive. A wise economic solution for this is the plant-microbial fuel cells in which a MFC is installed in the soil below plant roots, whose natural degradation causes the release of organic materials that can feed the bacteria^16^. Rather than feeding soil bacteria, roots also release redox species such as vitamins and NADPH that can donate electron at the anode to produce electricity^17,18^. The demand to provide food for the bacteria in MFCs can be overcome by utilization of photosynthetic micro-organisms such as cyanobacteria and microalgae that can synthesize their own carbon source^13,19–22^. Rather than using photosynthetic micro-organisms, bulk macro-organisms such as seaweeds and plant leaves may also be integrated in BECs^23–25^. Similar to bio-films which produce more electricity than singles bacterial cells^26^ photosynthetic native tissues such as plant leaves and seaweeds based BECs produce a more electricity than photosynthetic microbial fuel cells^21^. Another interesting approach for electricity generation is based on the integration of isolated components such as enzymes^27–31^, mitochondria^32^, chloroplasts^33^ and photosystems^34–37^ in BECs. An advantage of such methods is that the pure component which produces the electrical current is associated directly with the anode while the active redox species has less barriers to cross in order to reach the anode. Nevertheless, the purified component may lose it stability and activity faster than in living cells because of the lack of repair mechanisms or regeneration of the redox species^38^. Recent studies showed for the first time that BECs may also consist of the eye cell line *ARPE19*^39^ which produces an electric current that is greater than intact bacteria. The understanding that BECs are not limited to microorganisms and may utilize mammalian cell lines has opened a new unexplored avenue for studying the performance and electron transfer mechanism of new mammalian cell types that may be used for bio-electricity production. In this work, we produce current from 3T3 fibroblast cells in a BEC and apply electrochemical and spectroscopical methods to identify redox species released by the cells that may be involved in the external electron transfer between the inner area of the cells and the external anode. Also, we show that the cultivation media of the cells consists of redox species that can generate current in dark and under solar irradiation.

## Materials and methods

### Materials

All chemicals were purchased from Merck.

### 3T3 cell culture

NIH 3T3 fibroblasts were cultured in DMEM supplemented with 10% fetal bovine serum, nonessential amino acids, sodium pyruvate, L-glutamine, 100 U/ml penicillin, 100 μg/ml streptomycin, and 100 μg/ml neomycin. In all experiments, the 3T3 fibroblast cells were used in passage 7-15 and cultured in a 37°C humid incubator.

### Electrochemical measurement

All electrochemical measurements were performed using a Emstat4 conjugated with screen printed electrodes (SPEs) (Basi), with a graphite working and counter electrodes, and Ag coated with AgCl reference electrode. A drop of 70 μL of the analyte was placed on top of the SPE covering the entire area the electrodes. In the case of cells the measurements were started 5 min after the placement of the drop to enable the cells to sink from the drop suspension down to the electrodes. Photoelectrochemical measurements were done by placing a white light emitting diode (LED) above the SPEs. The light intensity at the SPEs surface height was measured buy a light meter and set to 1000W/m^2^ by adjusting the distance of the light source from the SPEs. Chronoamperometry was measured applying an electrical potential of 0.9 V on the working electrode. Cyclic Voltammetry was conducted sweeping the potential from 0 to 1.2 V with a scan rate of 0.1 V/s.

### Spectroscopic measurements

All spectroscopic measurements were conducted by Tecan plate reader in 96 wells microplates. The volume of the measures analyte inside the well was 100 μL. Absorption measurements were done in the wavelength range of 300 – 800 nm. In all Fluorescence measurements for NADH detection (λ_excitation_ = 350 nm, λ_emission_ = 400 – 700 nm), and for flavins (λ_excitation_ = 450 nm, λ_emission_ = 500 – 800 nm), were measured with a bandwidth of 20 nm with an integration time of 40 μs.

## Results and Discussion

### Analysis of redox-active species in the cultivation medium for cell culture

Previous work has reported that the bacterial cultivation media Luria-Broth (LB) which mostly consists of yeast extracts of redox species such as NADH and flavines can generate electrical and photoelectrical current when integrated with the anode of a BEC. We postulated that the cultivation media of mammalian cells in which Dulbecco′s Modified Eagle′s Medium (DMEM) and fetal bovine serum (FBS) are major ingredients may also consist of redox species that may be utilized for current production. To study this, we have conducted cyclic voltammetry (CV) measurements of DMEM, FBS, and cell’s cultivation media. Saline solution (0.9% NaCl) which does not consist of any redox-active species was measured as a control experiment (Fig. 1a-d). The CV measurements showed a small anodic peak around 0.6 V for DMEM, two anodic peaks around 0.6 and 0.9 V for FBS and the full composition of the cultivation media. A current increase without any peaks shape was obtained for the Saline between 0.5 to 1.2 V. This current increase (which was obtained in all measurements) originates from oxygen evolution reaction. The peaks around 0.6 and 0.9 V have the voltametric fingerprints of Tryptophan and NADH respectively. These components were previously detected in LB media. To strengthen the hypothesis that NADH exists in the cultivation media, we have conducted fluorescence measurements (λ_excitation_ = 350 nm, λ_emission_ = 400 – 700 nm) (Fig. 1e). The obtained spectra showed the spectral fingerprint of NADH in both FBS and the cultivation media in which the fluorescence intensity was higher.

**Fig. 1.**
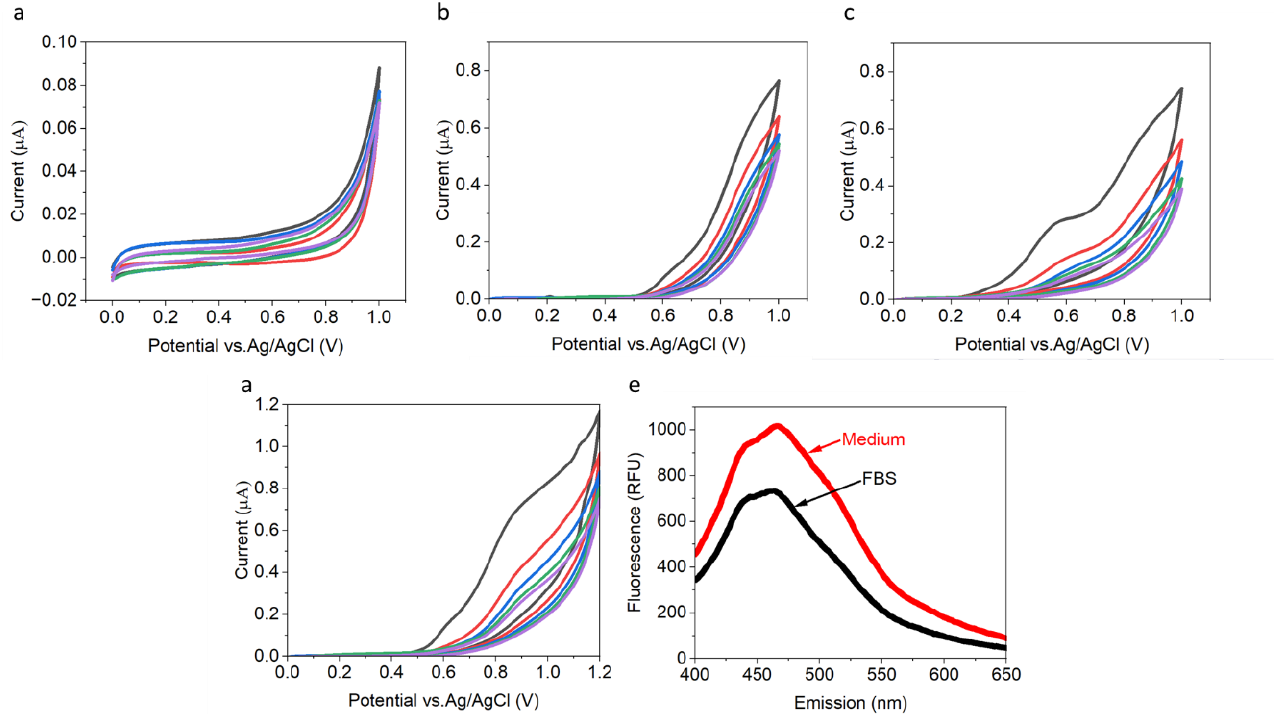
Analysis of redox-active species in the cell culture medium. CV and fluorescence spectra of the cultivation medium and its ingredients. **a** CV of Saline solution. **b** CV of DMEM. **c** CV of FBS. **d** CV of the cultivation medium. **e** Fluorescence spectra of FBS (black) and cultivation media (red) (λ_excitation_ = 350 nm, λ_emission_ = 400 – 700 nm). The excitation and emission wavelengths are compatible with the maximal fluorescence of NADH. In all CV voltammograms 5 cycles were measured. Cycles 1 – 5 are shown in black, red, blue, green, and purple respectively.

### Dark and Photocurrent production from cultivation medium and its ingredients

Previous work has reported the generation of dark and photocurrent from the bacterial cultivation medium LB, which was partially produced by NADH and flavins that originate from yeast extracts. As these molecules were also identified in the mammalian cells cultivation media, we wished to assess whether the media can generate electricity in dark and light while being integrated with an electrochemical cell. To do this, Chronoamperometry (CA) of Saline, DMEM, FBS and the complete cultivation media was measured using SPEs, with an applied potential of 0.9 V at the anode. This potential was chosen based on a peak at the highest potential obtained in the CV of the cultivation media (Fig. 1d). Irradiation of white light was conducted by a white LED from above (Fig.2a), with an intensity of 1000 W/m^2^ measured at the height of the SPEs. Light was turned on after 400s in dark. A decay of current for DMEM, FBS and the complete media but not for Saline was obtained over time. We sought that this decay is originating from a decrease in electron donors that are being consumed over time. The dark current of DMEM, FBS and the medium was ∼ 4.45, 5.65, and 6.79 μA / cm^2^ respectively. No photocurrent was obtained for the Saline and DMEM, and a maximal photocurrent of 0.6 and 1 μA / cm^2^ was obtained for FBS and the medium respectively (Fig. 2b). Based on the obtained results, we conclude that FBS plays a major role in the dark and photocurrent generation of the medium, while DMEM contributes to the dark current only.

**Fig. 2.**
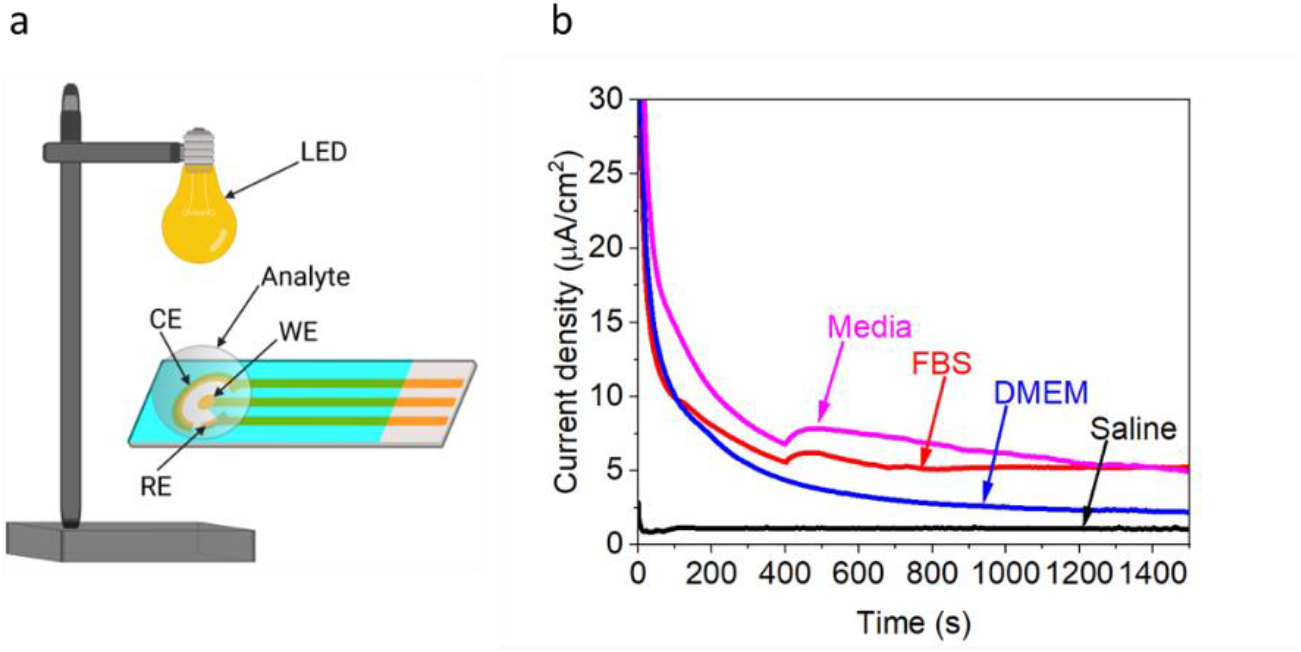
Dark and photocurrent production from cultivation medium and its ingredients. CA of Saline DMEM, FBS, and the complete cultivation media was measured in dark and under white light irradiation (1000 W/ m^2^) measured with an applied potential of 0.9 V on the anode.

### Fibroblast cells release NADH and flavines to the external cell solution

A previous work has reported for the first time the use of mammalian cells as electron donors when integrated in a BEC^39^ using *ARPE19* eyes cell line. The current production by these cells was partially generated by the release of NADH and flavins to the external cell solution (ECS) which can act as electron donors. To construct connective tissues fibroblast cells release numerous molecules to the ECS. While various proteins-based compounds secreted by fibroblast were previously reported, the identification of redox-species from these cells was not studied by far. To differentiate the molecules released by the cells from the ones exist in the media, the cells were centrifuged and their cultivation media was replaced with Saline. A drop of 70 μL of cells solution was placed on the SPE and incubated for 10 min to enable them to sink down to the electrodes surface. CV of the cells was measured to identify redox-activity (Fig.3a). The obtained voltammogram showed peaks around 0.6 V which correspond to the fingerprint of tryptophan suggesting that redox proteins may be released by the cell. An additional peak around 0.9 V corresponds to the fingerprint of NADH. Interestingly, similar peaks were previously reported for *ARPE19* cells^39^. To strengthen the suggestion that NADH is released by the cells and evaluate its quantity in comparison with the NADH pools inside the cells, Fluorescence spectra of the cells and the ECS were measured (λ_excitation_ = 350 nm, λ_emission_ = 400 – 700 nm) (Fig. 3b). The obtained spectra showed that the NADH quantities in the ECS is about 14 % from the internal content of the cells. We suggest that the NADH in the ECS may originate from molecules released by the cells and also the spilled content of dead cells.

**Fig. 3.**
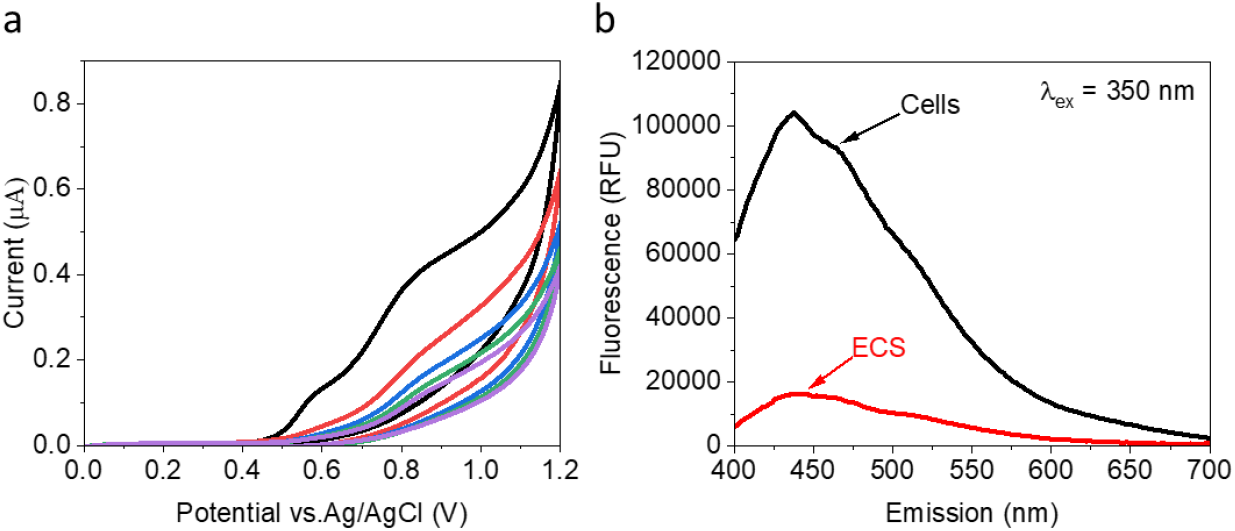
Fibroblast cells release NADH and flavines to the ECS. CV of the cells and fluorescence measurements of their ECS. **a** CV of fibroblast cells, (0 - 1.2 V, scan rate of 0.1 V/s). Cycles 1 – 5 are shown in black, red, blue, green, and purple respectively. **b** Fluorescence spectra of Fibroblast cells (black) and their ECS (red). The samples were excited at 350 nm, which is compatible with the maximal intensity of NADH.

### Live Fibroblast cells produce electrical current in a BEC

Following the identification of redox species released by fibroblast cells, we next wished to assess whether live fibroblasts can be utilized to generate electrical current in a BEC. For this purpose, CA of intact fibroblasts and their ECS was measured with an applied potential of 0.9 V at the anode (Fig. 4). This potential was evaluated to be optimal as it enables to cause oxidation of the redox species whose presence was observed in the CV between 0.5 to 0.9 V (Fig. 3a). The current produced by the ECS was decaying as function of time. We suggest that this decay derives from the consumption of electron donors such as NADH that are being oxidized at the anode. The CA curve of the cells started with an exponential decay between 0 – 400 s. We postulate that the sharp current decrease between 0 to 200 s originates from the capacity current while the small decrease between ∼ 200 to 400 s results from electron donors in the ECS accumulated prior to the beginning of the measurements. A small current increase of about 1 μA / cm^2^ was obtained around 400s which turned into a relatively constant current from 700 to 1500 s. We suggest that this current represent an equilibrium in which the rate of the electron donors release is almost equal to the rate of their consumption. The maximal current density between 400 to 1500 s was about 7.5 μA / cm^2^ which is similar to the current density previously obtained by *ARPE19* cells ^39^.

**Fig. 4.**
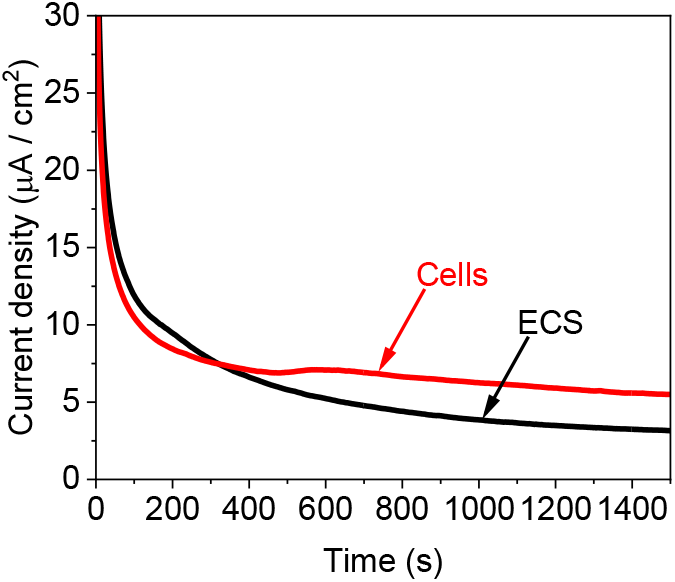
Live Fibroblast cells produce electrical current in a BEC. CA of ECS (Black) and Fibroblast cells (red). A potential of 0.9 V was applied on the anode.

## Conclusions

In this work, we showed that electrical dark and photo current can be generated by the cultivation medium of mammalian cells which consist of NADH and redox-proteins that may play a role in the electricity generation. Also, we show for the first time that Fibroblasts can be integrated in a BEC to generate current while NADH and redox-proteins may be the major electron mediator between the inside of the cells and the external anode. The concept of using fibroblast cells and the studying of their external electron transfer mechanisms may pave the way for future energy and biomedical technologies

## Acknowledgments

We thank Dr. Ines Zucker of Tel-Aviv University for the technical assistance in performing spectroscopic measurements.

## Author contributions

YS and AL conceived the idea. YS designed the experiments. YS performed the main experiments. YS, and AY wrote the paper. AY supervised the entire research project and provided funding.

